# An interactive mass spectrometry atlas of histone posttranslational modifications in T-cell acute leukemia

**DOI:** 10.1101/2022.05.05.490796

**Authors:** Lien Provez, Bart Van Puyvelde, Laura Corveleyn, Nina Demeulemeester, Sigrid Verhelst, Béatrice Lintermans, Simon Daled, Juliette Roels, Lieven Clement, Lennart Martens, Dieter Deforce, Pieter Van Vlierberghe, Maarten Dhaenens

## Abstract

The holistic nature of omics studies makes them ideally suited to generate hypotheses on health and disease. Sequencing-based genomics and mass spectrometry (MS)-based proteomics are linked through epigenetic regulation mechanisms. However, epigenomics is currently mainly focused on DNA methylation status using sequencing technologies, while studying histone posttranslational modifications (hPTMs) using MS is lagging, partly because reuse of raw data is impractical. Yet, targeting hPTMs using epidrugs is an established promising research avenue in cancer treatment. Therefore, we here present the most comprehensive MS-based preprocessed hPTM atlas to date, including 21 T-cell acute lymphoblastic leukemia (T-ALL) cell lines. We present the data in an intuitive and browsable single licensed Progenesis QIP project and provide all essential quality metrics, allowing users to assess the quality of the data, edit individual peptides, try novel annotation algorithms and export both peptide and protein data for downstream analyses, exemplified by the PeptidoformViz tool. This data resource sets the stage for generalizing MS-based histone analysis and provides the first reusable histone dataset for epidrug development.

## Background & Summary

Epigenomics studies the epigenetic changes that occur in organisms to modulate gene activity and induce phenotypic changes in response to the environment. Because epigenetic changes can lead to run-away cell states and cancer, inhibiting epigenetic mechanisms like DNA methylation and histone posttranslational modifications (hPTMs) is an appealing strategy to prevent and revert disease signatures. Recurrent alterations in genes that regulate epigenetic processes have taken hematopoietic malignancies and especially T-cell acute lymphoblastic leukemia (T-ALL) to center stage in the development of drugs that target such modifications, i.e. epidrugs. These epidrugs have been approved in multiple clinical trials and it is now clear that reprogramming the epigenetic landscape in the cancer epigenome holds great promise for patient care^1–4^. However, targeting of essential epigenetic mediators is being hampered because the mapping of hPTMs using mass spectrometry (MS) is inconvenienced by complex data analysis and impractical raw data reuse.

Therefore, we here present the very first preprocessed atlas of the histone code for 21 T-ALL cell lines, measured by MS. In essence, an LC-MS system measures the intensity and physicochemical properties of analytes like *m/z*, retention time (*t*_*R*_) and fragmentation pattern, turning a physical sample into a digital transcript. Yet, compared to mining sequencing data, extracting biological information from this raw data matrix is very challenging and is hypothesis-driven, especially for histones, given the combinatorial complexity of the histone code^5–7^. Moreover, the current way of data acquisition and sharing is not tailored to reuse, because (i) the datasets lack power because they are small, often limited by the nanoflow LC stability^8^, (ii) data quality metrics are not always provided, (iii) the selection of hPTMs included in the search is research-driven and (iv) histones require extensive manual data curation by an expert, because of the many (near-)isobaric peptidoforms and ambiguous annotations they create^9^. Therefore, a larger, quality controlled and preprocessed dataset can uniquely allow (bioinformatics) researchers to tailor the analysis of the data to their own hypotheses, helping to mediate the current reproducibility crisis in cancer research^10^.

The histone atlas presented here contains six biological replicates of 21 T-ALL cell lines, acquired in a single full factorial design, interspersed with a quality control sample made from all samples in the experiment, totaling 161 LCMS runs. The complete instrumental quality control report is freely accessible and editable at Skyline Panorama QC^11^. To assure the robustness of the measurements, eight selected cell lines were additionally grown and measured two years apart on a different instrument, revealing a reproducible abundance of peptidoforms and therefore the individual hPTMs over time, despite their dynamic nature. The most differential hPTM over these eight cell lines, H3K27me3, was confirmed with both western blot and flow cytometry, illustrating that MS provides a quantitative picture of hPTMs, without the need for targeted antibodies.

To enable data reuse, we have uniquely preprocessed the raw data by aligning all the measured precursor ions into a single, platform independent Progenesis QIP project that is fully licensed and thus fully functional (enabled in collaboration with Nonlinear Dynamics, Waters). As a reference, the ion population was annotated using Mascot with a selection of 9 hPTMs and all histone peptidoform precursor ions were manually curated by an expert (and flagged accordingly) to improve quantitative accuracy. Other (bioinformatics) users can now start with this intuitive and browsable template and i) create an experimental design containing their cell lines of interest, ii) export (certain) spectra for alternative searching strategies, iii) extend the manual feature curation, and iv) export the individual measurements to create quantitative reports and quantify specific hPTM targets not yet included in the current search. The latter is exemplified by an intuitive visualization tool (PeptidoformViz Shiny App) that can read exported *.csv files for downstream analyses, currently including advanced normalization strategies. Notably, the library also contains 505 proteins that were co-extracted, i.e. the acid extractome as described before^12^. The workflow of the experiment is given in **Figure 1**.

**Figure 1.**
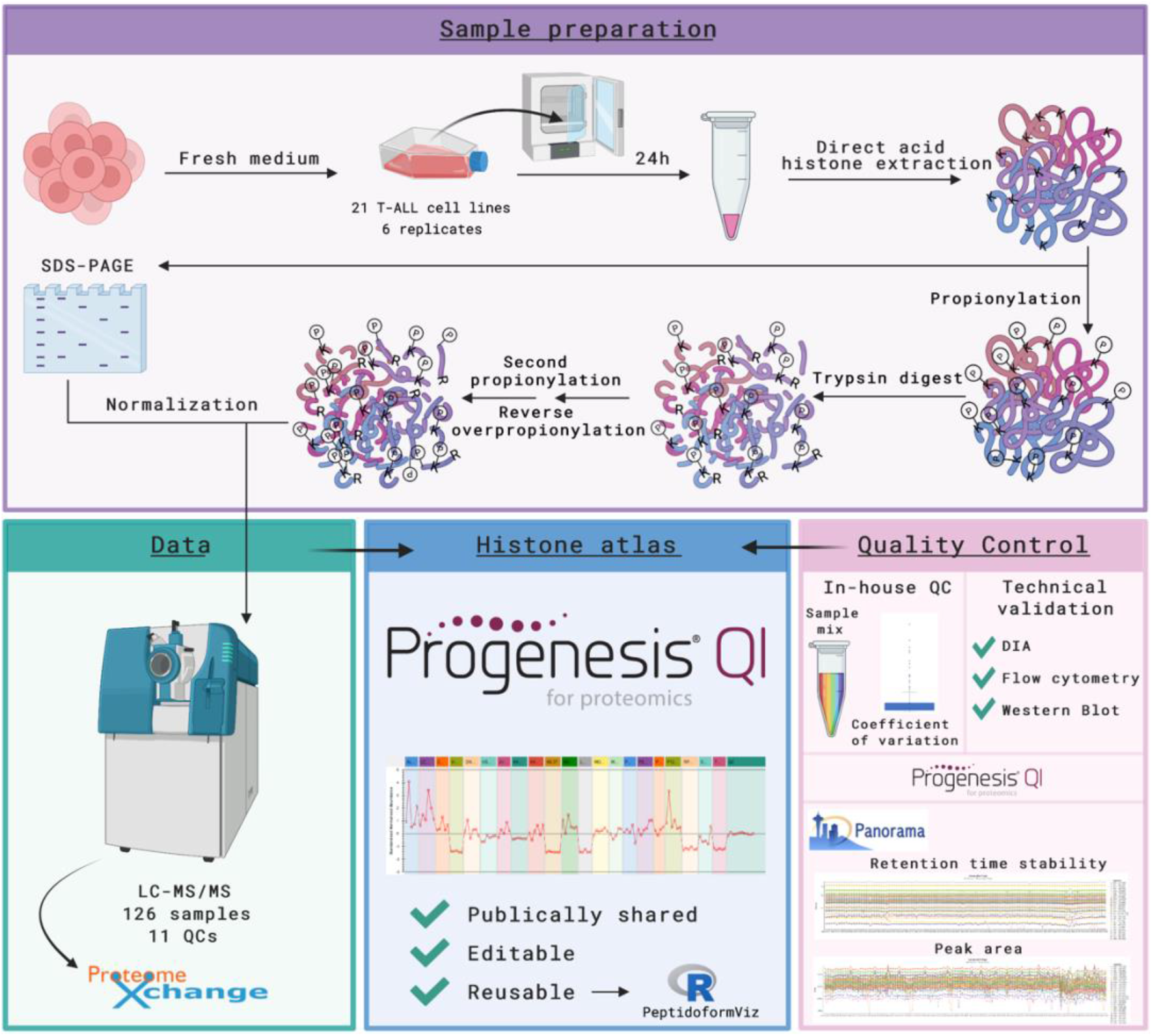
Overview of the study. T-ALL cells were collected 24 hours after medium was refreshed. After direct acid extraction, histones were propionylated and digested using trypsin to enable better separation on the mass spectrometer. Samples were measured using LC coupled to MS/MS; raw data is shared on ProteomeXchange. The experimental spectra were identified using Mascot and further processed in Progenesis QIP. The histone atlas is publicly shared as a Progenesis QIP project, which is editable and reusable a.e. by the PeptidoformViz tool. Multiple quality control steps were executed. Created with BioRender.com.

According to our knowledge, the size, format, and quality control of this atlas are unprecedented. We therefore express the hope that it will set the scene to not only help revolutionize epidrug development, but also contribute to the recent fields of toxicoepigenetics and pharmacoepigenetics, benefiting public health in a new way.

## Methods

### Cell culture and sample collection

T-cell acute lymphoblastic leukemia cell lines (DSMZ) were grown in culture flasks in RPMI 1640 medium (Gibco) supplemented with 10% (HSB-2, JURKAT, LOUCY, RPMI-8402, DND-41, KOPT-K1, PER-117, PF-382, SUP-T11) or 20% (ALL-SIL, HPB-ALL, MOLT-16, PEER, TALL-1, CCRF-CEM, CUTTL1, KARPAS 45 (JC), MOLT-4, KE-37, P12-ICHIKAWA) heat-inactivated fetal bovine serum (Sigma), penicillin (100 U/mL)-streptomycin (100 μg/mL) and 2 mM L-glutamin (Gibco) and incubated at 37°C with 5% CO_2_ and 95% humidity. Cultures were verified to be free of mycoplasma contamination using the TaKaRa PCR Mycoplasma Detection kit. Cells were incubated in fresh medium at a concentration of 1*10_6_ cells/mL for 24 hours before counting and harvesting cells by centrifugation (5 min, 1500 RPM, 4°C). After washing the cell pellets with 1 mL cold PBS, the cell pellets were snap-frozen in -150°C and stored in -80°C. For each cell line, 6 biological replicates were harvested from different subcultures at different time points.

### Histone extraction

Assessment of different extraction protocols described earlier, showed a higher histone yield with the direct acid (DA) extraction method compared to the hypotonic lysis buffer (HLB) method in a subset of T-ALL cell lines used in this study^13^. Before DA histone extraction, the snap-frozen cell pellets were resuspended in cold PBS and aliquoted in 1.5 mL Protein LoBind Eppendorf tubes to a concentration of 2*10^6^ cells per aliquot. Next, the pellets were centrifuged (10 min, 300 g, 4°C), resuspended in 250 µL 0.4 M HCl, and incubated for 2 hours on a rotator at 4°C to promote lysis of nuclei and solubilization of histones. After centrifugation (10 min, 16 000 x g, 4°C), the supernatant was transferred to a new Eppendorf tube. To promote precipitation of histones, 120 µL of 33% trichloroacetic acid was added slowly, after which the Eppendorf tubes were inverted several times and incubated on ice for 30 min. The pelleted histones (10 min, 16 000 x g, 4°C) were washed twice with slowly added ice-cold acetone (5 min, 16 000 x g, 4°C) and dried at room temperature. Pellets were resuspended in 150 µL milliQ water and centrifuged (16 000 × *g*, 10 min, 4°C) to remove the remaining insoluble pellet. 130 µL of the supernatant was transferred into fresh 0.5 mL Protein LoBind Eppendorfs for mass spectrometry and 20 µL was collected for gel electrophoresis. The histone extraction was performed on ice. Due to the high number of samples, samples were randomized into multiple batches for histone extraction (**Supplementary File 1**).

### Gel electrophoresis and image analysis

Extracted histones were vacuum-dried, after which 2x Laemmli-buffer with 2-mercaptoethanol (4% SDS, 20% glycerol, and 10% 2-mercaptoethanol in 50 mM Tris-HCl pH 6.8) was added. Next, samples were heated at 95°C for 10 minutes on a shaker to break the disulfide bonds and denature histones. Then, histones were separated at 100V and 250mA on 8-16% Criterion TGX precast gels (Bio-Rad Laboratories). After fixation with a buffer containing 7% acetic acid and 10% methanol, gels were stained with the fluorescent Sypro Ruby gel stain, scanned using a Versadoc imaging system (Bio-Rad Laboratories) and quantified with QuantityOne. On each gel, control lanes with a fixed amount of 2 µg bovine histones were included on each gel to facilitate quantification (**Repository file 1 on PXD031500**). In case no histones could be detected on gel or less than 5 µg histones were extracted, sample collection and histone extraction was repeated to ensure that 6 replicates for each cell line could be prepared for mass spectrometry (**Supplementary File 1**).

### Histone propionylation and digestion

In order to generate more hydrophobic peptides and enable better separation on the mass spectrometer, purified histones were propionylated as previously described^14^. Briefly, 5 µg of histones were vacuum dried and resuspended in 20 µL 1M triethylammonium bicarbonate buffer (TEABC). Next, 20 µL of propionylation reagent (propionic anhydride: 2-propanol 1:80 (v/v)) was added, the samples were vortexed and shaken at room temperature for exactly 30 minutes. During this first propionylation reaction the protein N-termini and the lysines become propionylated. To ensure that no aspecific propionylation occurs, the reaction was stopped by adding 20 µL MilliQ H_2_O, vortexing and shaking at 37°C for 30 minutes. Next, the samples were vacuum dried and digested overnight at 37°C using trypsin (at an enzyme/histone ratio of 1:20 (m/m)) in 500 mM TEABC, supplemented with CaCl_2_ and ACN to a final concentration of 1 mM and 5% respectively. After trypsin digestion, the samples were vacuum dried and a second propionylation was performed to propionylate the newly generated peptide N-termini. To reverse aspecific overpropionylation at serines, threonines and tyrosines, the samples were resuspended in 50 μL 0.5 M NH_2_OH and 15 μL NH_4_OH at pH 12 and incubated for 20 minutes at room temperature. Finally, 30 μL of 100% formic acid (FA) was added and samples were vacuum dried. Due to the extensive size of the sample batch, samples were randomized into multiple batches for propionylation (**Supplementary File 1**).

### Liquid chromatography and mass spectrometry acquisition

Liquid chromatography and mass spectrometry analysis was performed as previously described^12,15,16^. First, the propionylated histone peptides were resuspended in 0.1% FA in water (Biosolve). Next, a beta-galactosidase (SCIEX) and MPDS II (Waters Corporation) solution in 0.1% FA in water was added to ensure that an injection volume of 10 µL results in 2 µg histones, 62.5 fmol beta-galactosidase and 62.5 fmol MPDS II on column. Based on initial normalization factors, some samples were re-injected with adapted sample amounts to ensure an injection volume of 2 µg histones. A quality control (QC) sample was made by pooling 2 µL of each sample. Peptides were separated using a low pH reverse phase gradient on the NanoLC 425 system operating in capillary flow mode (5µL/min), coupled to a TripleTOF 6600+ mass spectrometer (SCIEX, Concord, Canada). The samples were loaded at 10µL/min with mobile phase A (0.1% FA in water) and trapped on a TriArt C18 guard column (YMC) (id 500µm, length 5mm, particle size 3 µm) for 3 minutes. A Phenomenex Luna Omega Polar C18 column (150 × 0.3 mm, particle size 3 µm) at 30°C was used at a flow rate of 5µL/min with a 60-minute gradient from 3-45% mobile phase B (0.1% FA in ACN), followed by a washing step at 80% B for 15 minutes. The sample list was randomized using a full factorial design and interspersed with QC injections. Each cycle of 2.3s consisted of one MS1 scan (m/z 350-1250) of 250ms, followed by MS2 data-dependent trigger events (m/z 100-1500, high sensitivity mode) of 200ms each. A maximum of 10 candidate precursor ions (charge state 2+ to 5+) exceeding 300 cps were monitored per cycle and these were then dynamically excluded for 10s. Ion source parameters were set to 4.5 kV for the ion spray voltage, 25 psi for the curtain gas, 10 psi for nebulizer gas (ion source gas 1), 20 psi for heater gas (ion source gas 2), and 100°C as source temperature.

### Data preprocessing

Since analysis of raw MS/MS data is very challenging, we therefore preprocessed the data to facilitate reuse, as previously described^15^. The workflow of the data processing is given in **Figure 2** and can now be adapted and executed with different parameters by other users. Briefly, raw data from all runs were imported and all runs were aligned in a single experiment in Progenesis QIP 4.2 (Nonlinear Dynamics, Waters) for feature detection. Next, the 20 MS/MS spectra closest to the elution apex were selected for each precursor ion and merged into a single *.mgf file to search in Mascot (Matrix Science). Mascot is a database search engine applies probabilistic scoring to define peptide-to-spectrum matches (PSMs). Two types of searches were performed on this file: 1) a quality search to verify the presence of non-propionylated standards (ß-gal) and to assess the extent of underpropionylation; and 2) an error tolerant search to identify the proteins present in the sample and to determine the best set of 9 hPTMs for further analysis. The Mascot search parameters used for both searches are shown in **Table 1**. A new FASTA-database was generated for further analysis based on the results from the error tolerant search^6^. Next, the three MS/MS spectra closest to the elution apex were merged into a single *.mgf and exported to perform a biological search in Mascot (**Table 1**). The search results (*.xml-format) were imported back into Progenesis QIP 4.2 to annotate the features from which they originated. Features that were annotated as histone peptidoforms were manually validated by an expert to resolve isobaric coeluting peptidoforms (**Figure 3**). Consequently, runs were normalized against all histone peptides to correct for differences in histone protein load across the samples. Note that for accurate hPTM quantification alternative normalization methods might be required a.e. in the PeptidoformViz tool. Outliers were removed based on normalization factor (greater than 10 and less then 0.3) and PCA clustering of all histone peptidoforms. Finally, deconvoluted peptide ion data (*.csv format) from all histones were exported from Progenesis QIP 4.2 for further statistical analysis.

**Table 1.** Overview of Mascot search parameters for the three different searches performed on this dataset.

| Parameter | Quality search | Error tolerant search | Biological search |
| --- | --- | --- | --- |
| Mass error precursor ions (ppm) | 10 | 10 | 10 |
| Mass error fragment ions (ppm) | 50 | 50 | 50 |
| Enzyme specificity | Trypsin | ArgC | ArgC |
| Miscleavages | 1 | 1 | 1 |
| Variable modifications | Propionylation (N-term)<br>Propionylation (K) | Deamidation (NQ)<br>Oxidation (M) | Acetylation (K)<br>Butyrylation (K)<br>Formylation (K)<br>Trimethylation (K)<br>Ubiquitination (K)<br>Methylation (R)<br>Dimethylation (KR)<br>Deamidation (NQR)<br>Phosphorylation (ST) |
| Fixed modifications | / | Propionylation (N-term)<br>Propionylation (K) | Propionylation (N-term)<br>Propionylation (K) |
| Database | Human Swissprot supplemented with contaminants from the common Repository for Adventitious Proteins (cRAP) database | Human Swissprot supplemented with contaminants from the common Repository for Adventitious Proteins (cRAP) database | Custom-made FASTA-database |

**Figure 2.**
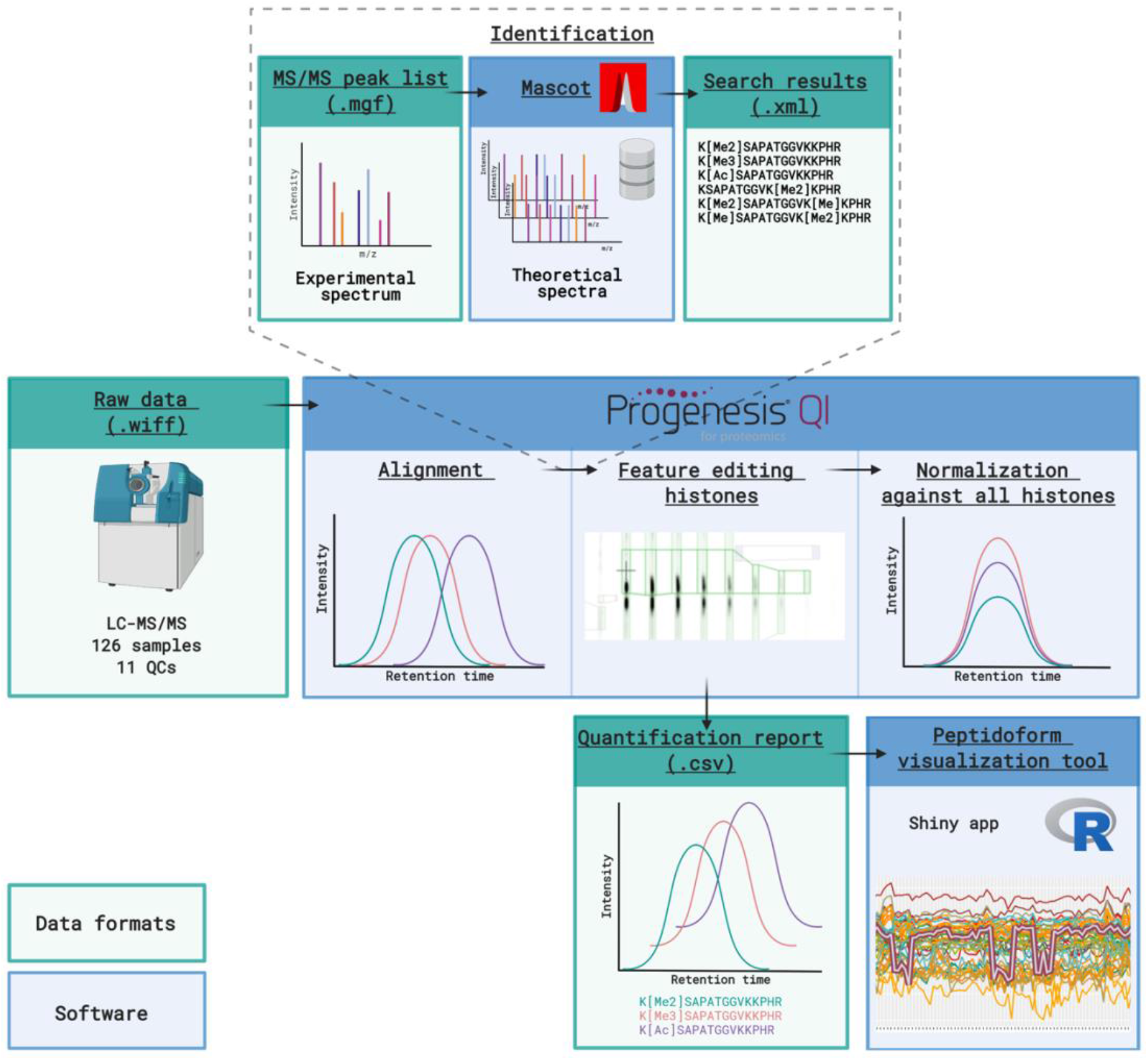
Workflow of the data processing. Raw MS/MS data is imported into Progenesis QIP and runs are automatically aligned using a QC sample as a reference. Next, MS/MS peak lists are exported from Progenesis QIP to perform peptide searches against a database of theoretical spectra in Mascot. Search results are imported back into Progenesis QIP and features identified as histones are manually edited. Then, normalization is performed against all histones to correct for differences in histone protein load across the samples. However, deconvoluted non-normalized peptide ion data from all histones are exported for normalization and visualization in the PeptidoformViz tool. Figure created with BioRender.com.

**Figure 3.**
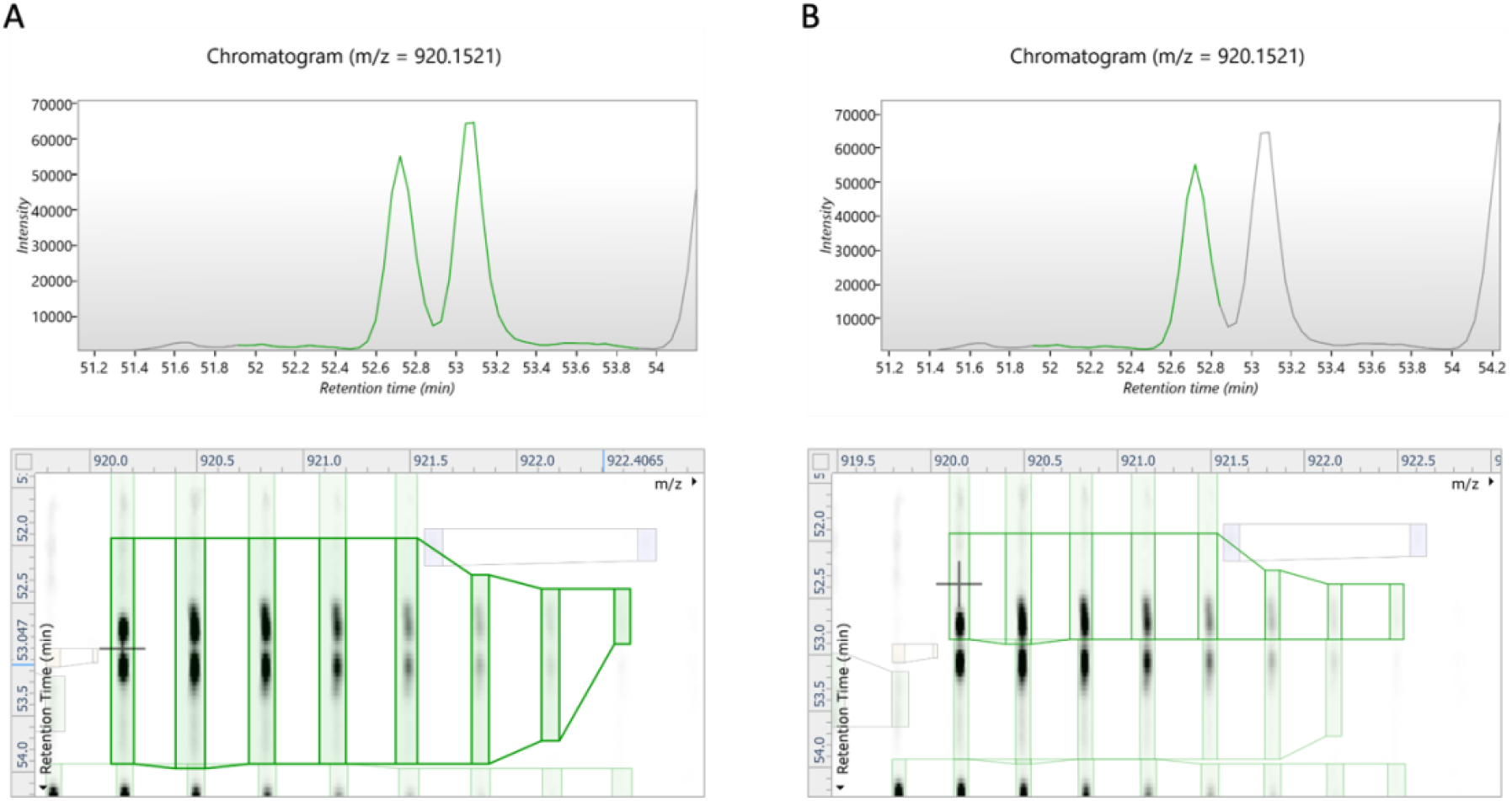
Example of a manual curated histone peptide in Progenesis QIP. **A**.Before feature editing, two baseline-separated isobaric peptide ions were merged as one feature. **B**. After feature editing, the two ions are now split into two separate features, allowing more accurate identification and quantification.

## Data Records

The mass spectrometry raw data (*.wiff and *.scan files generated by the instrument), data analysis files and repository files have been deposited to the ProteomeXchange Consortium (http://www.proteomexchange.org) via the Panorama partner repository with the dataset identifier PXD031500^17,18^. In addition, a Sample and Data Relationship File (SDRF)^19^, an Investigation Description File (IDF) and the sample list have been uploaded to ProteomeXchange, to better accommodate reanalysis of this comprehensive T-ALL hPTM dataset. Finally, the atlas is provided as a single archived Progenesis QIP project (*.ProgenesisQIPArchive) with embedded license, allowing everyone interested to browse through and further manually curate the data. Of note, the ProgenesisQIP software can be downloaded freely from the Nonlinear Dynamics website (https://www.nonlinear.com/progenesis/qi-for-proteomics/). The peptide quantification reports (*.csv) exported from Progenesis QIP, together with the metadata file can be used within the PeptidoformViz tool.

### Technical Validation

#### Sample preparation and data acquisition

Due to the considerable number of samples in the experiment, sample preparation was performed in several batches. However, this was done in a randomized fashion to minimize sample variation (**Supplementary File 1**). Acquisition was done according to the complete randomized block principle, i.e. each block being a sequence of one replicate of each condition (cell line). The sequence of measurement can be verified by the sample list uploaded to Panorama QC^11^. Every ten samples, 50 fmol of betagalactosidase (5600) or 40 fmol of PepCal Mix (6600+) standard was injected to calibrate the LC-MS instrument. Additionally, a mixture of all samples and MPDS and beta-galactosidase standards was made as a quality control (QC) to assess instrument performance throughout the batch.

#### Instrument performance

We performed a system suitability test to monitor LC-MS performance in a longitudinal fashion^11^. Therefore, we used the vendor neutral Panorama AutoQC framework and monitored signal intensity, retention time and mass accuracy. The set of peptides used to calculate these metrics were selected by searching the *.MGF file exported from Progenesis QIP with Mascot Daemon (v2.7) using a custom-made database of beta-galactosidase and the four proteins present in the MassPrep protein digestion standard mixture (MPDS2). From these five proteins, a total of 56 precursors was selected and subsequently appended to the Skyline target list. Using the AutoQC Loader software (version 21.1.0.158) a configuration file was set-up that enables the automatic import of each data file acquired with the SCIEX TripleTOF 6600+. The data and skyline reports were published to the PanoramaWeb folder “U of Ghent Pharma Biotech Lab – hPTM T-ALL Atlas”. As an illustration, we assessed the mass accuracy (expressed in ppm) as depicted in **Figure 4A**. Based on this Levey-Jennings plot, the MS instrument behaved very robustly and the instrument calibration runs with the pepcal peptide mixture can clearly be localized. Next to mass accuracy, we also confirmed that no significant drifts in retention time or potential problems with the LC system could be detected.

**Figure 4.**
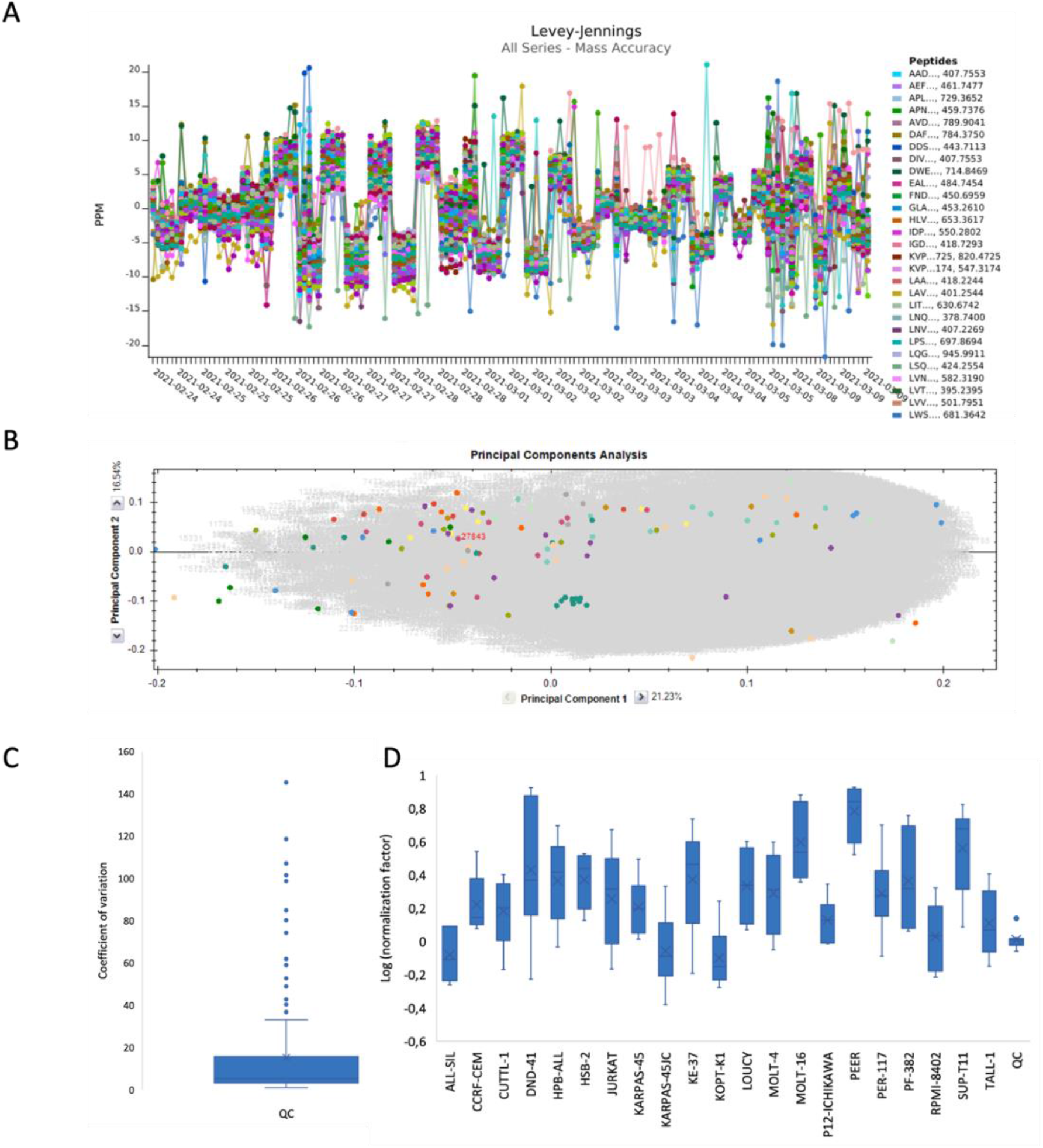
Metrics supporting the technical quality of instrument performance. **A**. Levey-Jennings plot of the mass accuracy for 56 selected precursors originating from the spiked-in protein digests of Beta-galactosidase and the MPDS2 mixture, highlights instrument performance throughout the batch. Mass calibrations were regularly performed, as seen by the repetitive leaps within +-10ppm **B**. Principal component analysis (PCA) in Progenesis QIP shows that all QC’s are clustered in the centre of the experiment. **C**. Boxplot of the coefficients of variation based on the normalized abundances of all histone features of the QC runs. **D**. Boxplot of the normalization factors for each cell line after outlier removal, based on normalization against all histones using a QC sample as a normalizing reference. Normalization factors are log transformed.

In Progenesis QIP, a principal component analysis (PCA) can be performed showing that all QC’s are clustered in the centre of the experiment (**Figure 4B**), thereby illustrating that the variations in precursor intensities caused by the measurement are negligible compared to the biological and sample preparation variation. Indeed, the coefficient of variation was calculated based on the normalized abundances of all histone features of the QC runs, providing an average CV of 15% in instrumental variation (**Repository file 2 on PXD031500, Figure 4C)**. Additionally, a separate QC metrics tab in the software allows to visualize sample preparation metrics, instrument metrics and experiment metrics. This is a complementary perspective to the QC metrics available in PanoramaQC. **Figure 4D** shows the normalization factors for all cell lines separately after outlier removal. Normalization was performed against all histones, using a QC sample as a normalizing reference.

#### Technical and biological robustness of the results

With two years apart, this experiment was conducted as a proof of concept (PoC) on a subset of 8 T-ALL cell lines. The exact same workflow was performed as described in the Methods section, except for MS data acquisition, which was carried out on a TripleTOF 5600 instrument (Sciex) in two different acquisition schemes. Data processing and analysis of DDA data were done using Progenesis QIP. Raw data and the archived Progenesis project can be found on ProteomeXchange with PXD identifier PXD031500 (archived with licenses). Data analysis of SWATH data was performed as published before^9^. To assess the technical and biological robustness of the results, a comparative analysis was performed between this batch and the same eight cell lines measured for the full atlas of 21 T-cell lines. Fold changes relative to the QC of a list of hPTMs of H3 and H4 were calculated for both batches and a multidimensional scaling (MDS) plot (**Figure 5A**) as well as a clustered heatmap (**Figure 5B**) were made in R (**Repository file 3 on PXD031500**). These indicate a successful biological repetition, adding value to the quality of the generated data.

**Figure 5.**
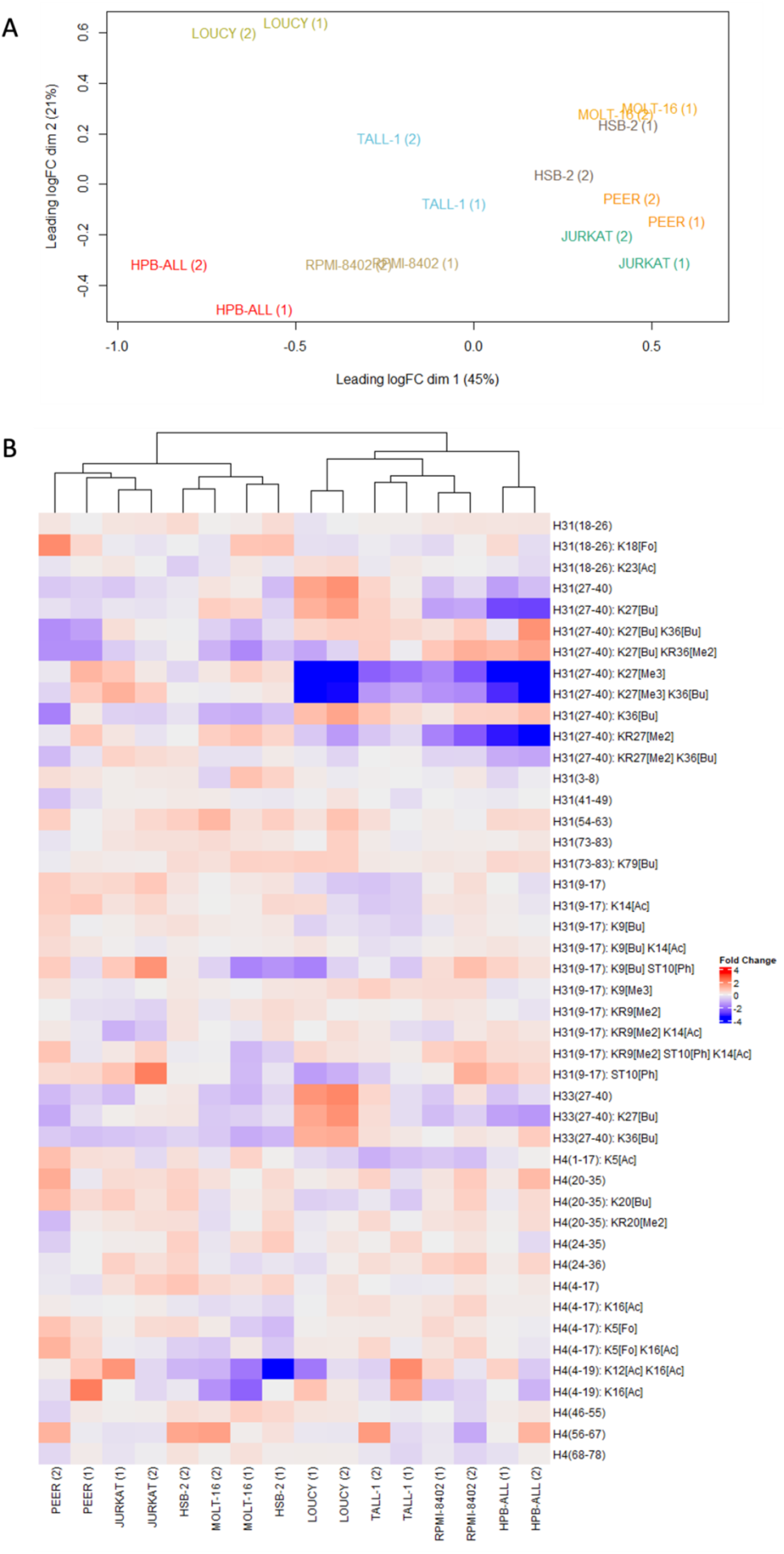
Technical and biological robustness of the data in two different batches. With two years apart, the exact same workflow was conducted on a subset of 8 T-ALL cell lines; however, data was acquired on a different instrument. Here, we show the robustness of the two different batches (1) and (2) **A**. The MDS plot shows clear clustering of the 8 cell lines over the two batches, expressed in fold changes relative to the QC. **B**. Clustered heatmap showing the fold changes relative to the QC for common peptidoforms of H3 and H4 for these 8 T-ALL cell lines measured in two different batches. After hierarchical clustering, all but one (HSB-2) cell lines clustered tightly together. Overall, this technical validation illustrates the robustness of the data.

In addition, we cross-validated the MS results obtained with DDA with three other analytical techniques. For the same 8 T-ALL cell lines used for the PoC above, the level of one of the most differential PTMs, namely H3K27me3, was investigated using sequential window acquisition of all theoretical fragment ion spectra (SWATH), western blot and flow cytometry (**Supplementary File 2**). All four techniques provided a similar pattern of H3K27me3 levels (**Figure 6**), indicating that mass spectrometry is a robust method to investigate the level of hPTMs in T-ALL. This divers level of H3K27me3 was within expectations since mutations and deletions within the PRC2 complex, responsible for the H3K27me3 modification, are frequently reported in T-ALL^20,21^. It has also been show that deregulation of this PRC2 complex can induce T-ALL^22^. Of note, observed hPTM levels might also be influenced by the ploidy state of the cell lines (**Supplementary File 3**).

**Figure 6.**
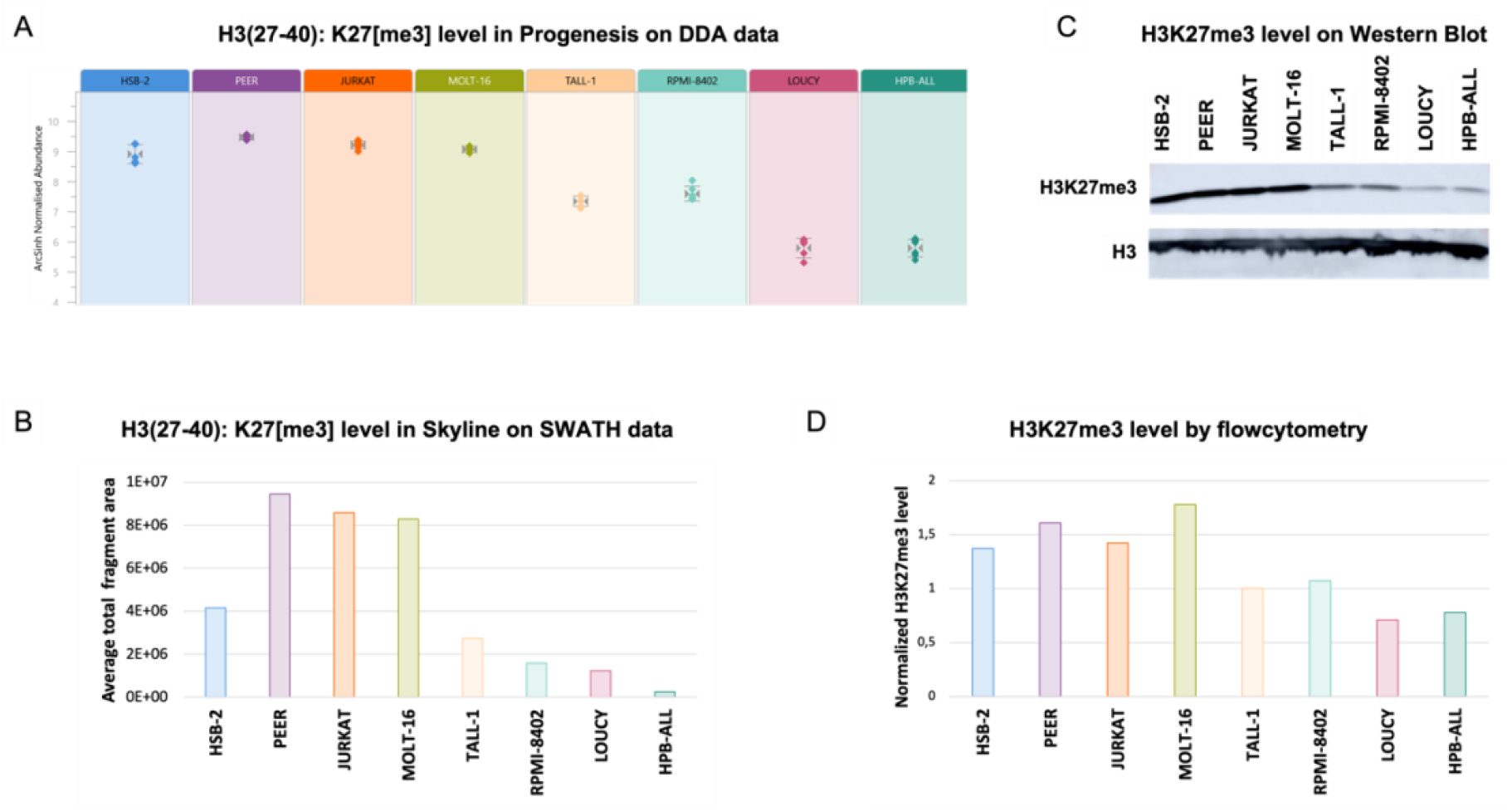
Cross validation of the H3K27me3 level. The level of H3K27me3, the most differential PTM based on DDA, in 8 cell lines was successfully verified with four different techniques. **A**. Normalized abundance of H3(27-40): K27[me3] obtained after analysis of DDA data in Progenesis QIP. **B**. The H3(27-40): K27[me3] level obtained after analysis of SWATH data in Skyline. The total level is obtained by summarizing the areas of the fragment ions. **C**. Western Blot analysis of H3K27me3. H3 was used as a loading control. **D**. H3K27me3 level measured by flow cytometry. The level was normalized against the H3 level.

### Usage Notes

This dataset provides the largest histone atlas to date for T-ALL and creates a framework for all research related to epidrug development for T-cell leukemia. One of the future perspectives of our own research group is to investigate the potential of hPTM levels as predictive biomarkers for epidrug outcome in T-ALL, i.e. pharmacoepigenetics. These potential biomarkers will be validated both *in vitro* and *in vivo*. Further optimization of MS analysis time could facilitate the analysis of patient samples in a clinical setting in the future^23^. Here we describe the functionalities available in the atlas, as an illustration of data reuse potential.

#### Data analysis

The raw data provided here can be processed in alternative ways. For example, raw data can be processed using a peak picking software of your choice that supports SCIEX data (*.wiff) (e.g. MSConvert), peptides can be annotated using alternative DDA data search engines (e.g. through SearchGUI) and further statistical analysis can be performed using alternative solutions. However, we here propose analyzing the preprocessed raw data using Progenesis QI for proteomics, which provides a straightforward data analysis workflow for DDA data with visualization at every stage. Progenesis QIP can be downloaded via https://www.nonlinear.com/progenesis/qi-for-proteomics/download/. Until recently, analysis in Progenesis QIP was only possible using either a hardware dongle or by entering license codes when prompted by the software. Hardware dongles are usually provided with a one-time purchase of the software and permanently allow the user to analyze all your data. Whereas with license codes, the users are limited to the analysis of an explicit number of runs. Once these are analyzed, the license code can no longer be used to enable any other runs. However, in agreement with Nonlinear Dynamics (Waters) we were able to publicly license our QIP project. As from now, this will be a feature that can be used for all QIP projects shared through ProteomeXchange. When opening the project in Progenesis QIP, the user will be able to adapt the entire data analysis workflow, for the runs already present in the project. A detailed user guide for DDA data can be found on the Nonlinear website (https://www.nonlinear.com/progenesis/qi-for-proteomics/v4.2/user-guide/) and a step-by-step video is available via https://www.youtube.com/watch?v=Mdrey1gauls.

An overview of the most prominent features in a Progenesis QIP experiment are shown in **Figure 7**. Briefly, after importing the raw data, Progenesis asks to automatically process the data, which includes automatic alignment and peak picking. Ions are aligned to compensate for a drift in retention time between runs. In the current project, Progenesis was allowed to automatically choose the most suitable alignment reference from all the QC runs (pooled samples). The next stage, ‘Review Alignment’, displays various graphics that allow the user to review the quality of alignment and edit it accordingly (**Figure 7B**).

**Figure 7.**
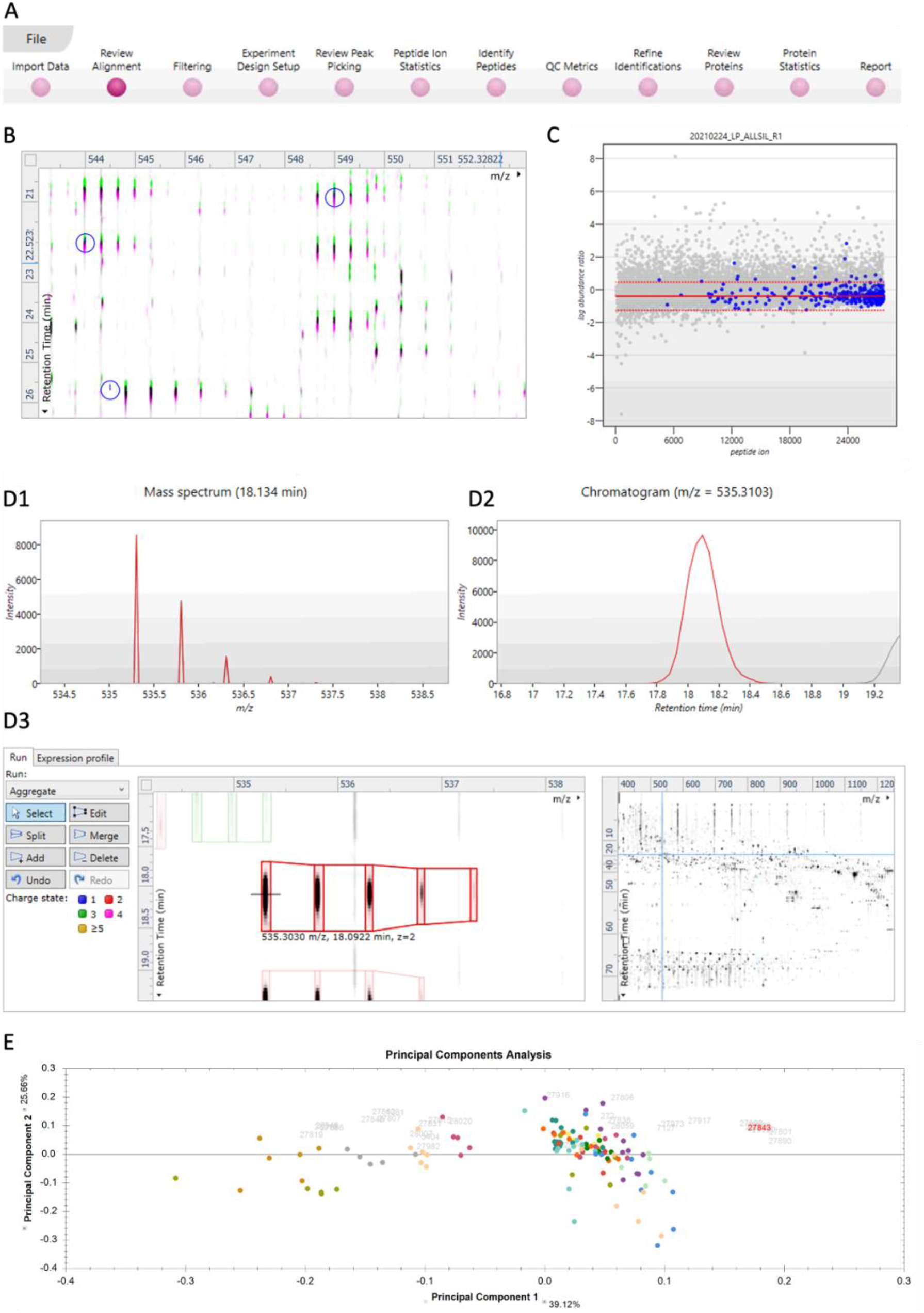
Overview of the most prominent features in a Progenesis QIP project. **A**. The data analysis pipeline as shown on the top of the screen in Progenesis. **B**. Vector editing on the ion intensity map. The current run is displayed in green and the reference run is displayed in pink. This window allows the user to review the alignment vectors in detail and manually add alignment vectors if required. **C**. Example of a normalization graph for one LC-MS run, as shown in the Filtering stage. Each blue dot shows the log of the abundance ratio for a different peptide ion. Peptide ions are shown ordered by ascending mean abundance. The normalization factor (solid red line) is then calculated by finding the mean of the log abundance ratios of the peptide ions that fall within the ‘robust estimated limits’ (dotted red lines). Peptide ions outside these limits are outliers and will therefore not affect normalization. **D**. The Review Peak Picking stage showing different data displays. **D1**. Mass spectrum for the current peptide ion in the selected run. **D2**. Chromatogram for the current peptide ion in the selected run. **D3**. Interactive 2D plot of the selected run, allowing the user to edit, split, merge, or delete current features and add new features. **E**. Principal Component Analysis (PCA) for the peptide stretch H3(27-40) at the Peptide Ion Statistics stage. Each solid-colored dot represents each run, with runs from the same condition, i.e., cell line, having the same color.

In the ‘Filtering tab’, any precursor ion that needs to be excluded for further analysis can be removed based on i) position (retention time and m/z), ii) charge state, iii) number of isotopes or iv) a combination of these ion properties. In this Progenesis QIP experiment, no filter was applied and thus all ions were used to perform normalization. Note that normalization will be updated if certain ion populations are removed, and the ‘Review normalization’ section will open automatically showing normalization plots for the remaining ions in each run. Progenesis will automatically select one of the runs that is ‘least different’ from all the other runs in the data set to use as a ‘Normalizing reference’. Then, a normalization factor is calculated for each sample to normalize it back to its reference (**Figure 7C**). The user can choose to i) normalize against all proteins, ii) normalize against a set of housekeeping proteins (e.g., histones), or iii) to not perform normalization.

In the next stage, different experimental designs can be created from the data. There are two main types of design that can be used: a between-subject design or a within-subject design. For this dataset, the samples of a given subject (i.e. T-ALL cell line) only appear in one condition (baseline) and thus a between-subject design was created to compare hPTM patterns between the 21 cell lines. However, one can choose to compare only a subset of cell lines by creating a new design. During further analysis, the user can switch between different designs by using the drop-down menu on the bottom left of the screen.

The next section, ‘Review Peak Picking’, allows the user to review and edit peptide ions of interest using different visual displays, i.e., 1D, 2D and 3D displays, which are shown in **Figure 7D**. Existing features can be edited, split, merged or deleted and new features can be added. However, in our current workflow, this section is only used to edit features that are annotated as histone peptides, ergo, identification of peptide ions is performed first. By right clicking on a peptide in the table, peptides with a certain property can be tagged to allow filtering in further analysis. The user can create their own tags, e.g. sort the peptide ions by abundance and highlight those that are high abundant in a certain cell line, or the user can choose one of the quick tags that are suggested by the software, i.e. tag according to i) Anova p-value, ii) max fold change, iii) certain modifications, iv) no MS/MS data or v) no protein ID. In the current project, tags were made for each histone protein and all tryptic peptides of H3 and H4, with the latter stated as e.g. H3(27-40). These tags can be used in the next stage, ‘Peptide Ion Statistics’, to perform different multivariate statistical analyses on the selected peptide ions. First, a principal component analysis (PCA) can be performed to verify if there are any outliers in the data and to check if the data clusters according to the user’s experimental design, as shown in **Figure 7E**. Second, a correlation analysis can be performed to evaluate the similarity in peptide expression between the different sample groups, which results in an interactive dendrogram. By clicking on a node in the branch, peptides with a certain expression profile will be highlighted (e.g., upregulated in LOUCY), which can then again be tagged.

In the next stage, ‘Identify Peptides’, the software allows the user to export MS/MS peak lists in various formats (e.g. mascot generic format) which can then be used by their respective search engines to perform peptide searches, since Progenesis QIP does not perform database searching for DDA data. By default, all available MS/MS spectra in the user’s experiment will be exported, of which the number is displayed on the export button. However, this set of MS/MS spectra can be filtered using the custom-made tags (e.g., peptide ions that are upregulated in a certain cell line of interest) and can be further refined based on quantity and quality of the spectra. In order to choose which spectra the user wishes to include or exclude from their export, the export button needs to be ticked in the table and then the ‘Batch inclusion criteria’ needs to be expanded, e.g., we used “Rank less than 4” to only include the three MSMS spectra closest to the elution apex of each peptide ion. When the user makes his selection, select the appropriate search engine, e.g., Mascot, and click ‘Export spectra’ to save a merged file. As mentioned before, due to the combinatorial explosion of hPTMs, the number of variable modifications that can be searched in Mascot is limited to 9. However, multiple identification files obtained from sequentially searching the MS/MS peak list with different search parameters (e.g. combinations of hPTMs) can be imported in the experiment. Note that for clarity, the hPTM abbreviations were prepared before importing the Mascot xml files back into Progenesis.

In order to investigate the quality of the data in the experiment, go to the next stage in the workflow, ‘QC metrics’. Here, the user can explore i) sample preparation metrics, i.e., displaying issues with sample preparation, ii) instrument metrics, i.e., showing whether the LC-MS system is performing robustly and iii) experiment metrics, i.e., highlighting any outliers in term of identified proteins and peptides in the experiment. This allows the user to examine whether there are any systematic effects throughout the processing that may affect the conclusion. Metrics of interest or concern can be flagged and exported into a report.

In the next stage, ‘Refine identifications’, unwanted or irrelevant peptide results can be removed based on several criteria i.e., score, hits, mass error, etc. Next, in the ‘Review proteins’ section, all details of the data are shown at the protein level, which allows the user to tag certain proteins of interest, e.g. to investigate histone variant expression across the 21 cell lines. Additionally, histone extraction by direct acid extraction leads to co-extracted proteins, the so-called acid extractome, which was proven to contain very useful biological information^12^. At this stage, the software also enables exporting the data to external pathway tools or export them directly as a *.csv file to continue further analysis. When interested in differential abundance analysis, an R-package like msqrob2 is highly recommended, as it enables quantitative peptidoform and protein-level statistical inference on LC-MS proteomics data (https://www.bioconductor.org/packages/release/bioc/html/msqrob2.html)^24^.

Moving on to the next stage, ‘Protein statistics’, comparable to Peptide statistics earlier in the workflow, Progenesis facilitates multivariate statistical analysis at the protein level. Finally, in the ‘Report stage’ a report of the experiment at the protein or peptide level can be exported as *.html file.

#### Visualizing the peptidoform level data

We here propose to export a *.csv file from Progenesis QIP containing deconvoluted non-normalized peptide ion data that can be imported into a newly made shiny app to visualize peptidoform level data. It facilitates the import, preprocessing and visualization of peptide level intensity data and will be updated to include advanced statistical analysis in future versions. This tool, called PeptidoformViz, is an interactive web app developed with the Shiny R Package (https://shiny.rstudio.com/, version 1.7.1) and can be downloaded from: https://github.com/statOmics/PeptidoformViz. Of note, an installation of R and RStudio is required^25,26^. Further installation instructions are provided on the GitHub webpage as linked above.

The app currently consists of three main panels: data import, preprocessing and visualization. The data import tab requires a tab, comma, space or semicolon separated file that contains peptide level intensity data. It should be structured so that each peptidoform occupies one row in the document and should include, on the column level, information on the protein ID, peptide sequence, modifications per peptidoform and intensities. If multiple samples were run in the same experiment, each sample should have its own intensity column. The intensity columns should be named in a way that one string pattern can be matched to all (e.g., all intensity columns start with the pattern “Intensity_”). E.g.:

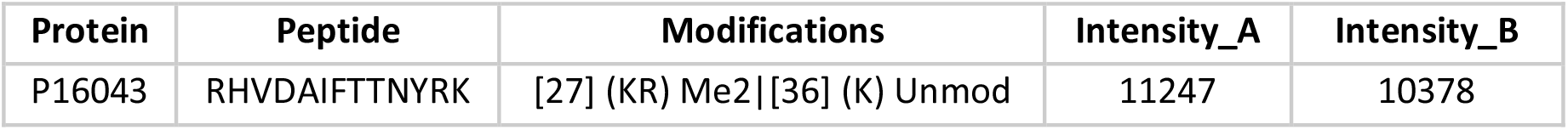

The import tab also requires a metadata file that contains the names of all intensity columns from the peptide level data file in one column and in the other columns identifying information ((e.g. different conditions such as species, treatment, cell line). These columns should contain at least enough information so that each row represents one unique ID. E.g.:

| Intensity columns | Sample | Condition |
| --- | --- | --- |
| Intensity_A | 1 | A |
| Intensity_B | 2 | B |

Alternatively, the user has the option to read in an example dataset via the “Read in example data button”. In this case, no other information has to be provided in the import tab. The example dataset included in the shiny app is in fact the dataset presented in this paper. In this way, it is readily accessible in the app and users can immediately start exploring it.

In the preprocessing panel, the user can opt for several preprocessing steps: logarithmic transformation, normalization and a filtering step based on missing intensity values. A density plot (**Figure 8 A1**) and boxplot (**Figure 8 B1**) of the data are generated to show the user the impact of the preprocessing steps. Intensities are usually lognormally distributed, hence, we recommend always including this transformation, unless the data have been log transformed before uploading.

**Figure 8.**
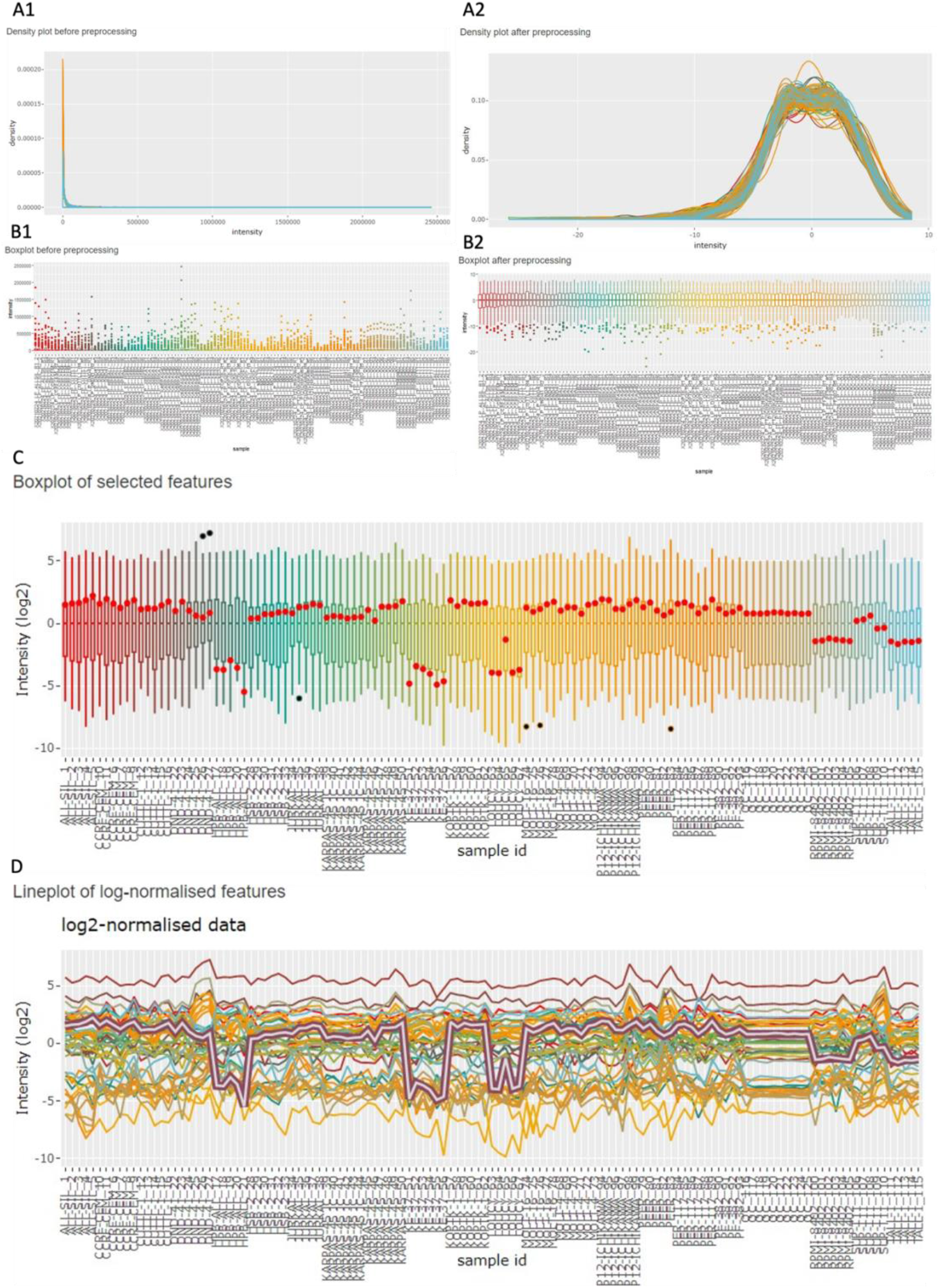
Data processing and visualization in the PeptidoformViz Tool. A and B can be found under the preprocessing tab, plots C and D under the visualization tab in the PeptidoformViz tool. **A**. The density plot both before (**A1**) and after (**A2**) preprocessing. The density plot visualizes the distribution of all peptidoform intensities for all proteins in each sample. In this example, this distribution is highly right skewed before preprocessing. **B**. The boxplot both before (**B1**) and after (**B2**) preprocessing. The boxplots visualize the distribution of the intensity values per sample (the different samples are present in the x-axis). The right skew before preprocessing can also be seen here: a lot of outliers are present above the third quartile. Upon logarithmic transformation and median centering, the boxplots reveal less outliers, as well as a median of zero for each sample. **C**. Boxplot in which each box represents the distribution of intensity values from the chosen protein in the corresponding sample, which are found on the x-axis. If one or more peptidoforms were highlighted, they will be displayed as red dots **D**. Line plot: each line represents a peptidoform from the data table. The y-axis represents the intensity, on the x-axis, the different samples are shown, making it easy to see in which samples (e.g., with a certain treatment, or from a particular cell line) the peptidoform shows a different intensity pattern. The selected peptidoform, in this example H3(27-40): K27[me3], is shown as a purple encompassed grey line in the line plot and as red dots in the boxplot.

However, do keep in mind that the intensities will then be displayed on the logarithmic scale. Users can choose between a logarithmic transformation with base 2, 10 or *e*. Furthermore, a mean centering or median centering normalization can be performed to normalize the marginal peptidoform distributions across samples, i.e. by subtracting the sample mean or median peptidoform intensity from all peptidoform intensities from a sample. This approach can help reduce systematic technical variation^27^. When the dataset contains many missing intensity values, it can be useful to filter out peptidoforms that were only identified/quantified in few samples. In our tool, the user can choose in how many samples each peptidoform should be at least observed (defaults to at least two). In this tab, general information (“stats”) about the number of proteins, peptides and peptidoforms is also displayed to inform the user on how the filtering step affects the data that will be used downstream. When all options are filled out, the user can click the preprocess button to perform all preprocessing steps and to generate the density plot (**Figure 8 A2**), boxplot (**Figure 8 B2**) and data stats upon preprocessing. Do note that this step should be completed before the user can go on to the visualization tab.

Upon preprocessing, the user can go on to the visualization panel. This panel consists of a protein selection and a normalization step, a data table, a line plot and a boxplot. Here we work with the peptidoforms of one protein only. Accordingly, the user can select a protein of interest present in the dataset via a dropdown menu. In the preprocessing step, users have the choice to perform a global normalization, i.e., based on all intensities for all peptidoforms in a sample. In the data visualization tab, users again have the option to normalize the peptidoform intensities towards the average/median peptidoform intensity of the selected protein in the sample. Hence, the user can choose to visualize absolute peptidoform abundances or relative abundances. The latter are useful to visualize peptidoform usage/switches that are not triggered by an overall change in the protein abundance. In the visualization panel, a data table is displayed containing all peptidoforms of the selected protein and their corresponding intensity values on the original scale or the log scale depending on users choice in the preprocessing tab. It is possible to search for certain patterns in peptide sequences in the table, e.g., a particular modification or a certain sequence. The data table can also be downloaded to the user’s hard disk. Two figures are rendered based on the displayed data table: a boxplot (**Figure 8C**) and a line plot (**Figure 8D**) of all the peptidoforms of the protein. In the boxplot, each separate boxplot corresponds to all intensity values of the selected protein in a particular sample. Each line in the lineplot corresponds to one peptidoform and displays the intensity value of a specific peptidoform over the different samples. In this way, the user can spot patterns in the (relative) peptidoform intensities across samples, e.g., a sharp change in intensity for treated samples vs non-treated samples. It is possible to highlight peptidoforms of interest by clicking on them in the data table. The corresponding line(s) in the lineplot will then be highlighted, and corresponding dots will appear in the boxplot. Both plots are interactive, so the user can zoom and pan on the figure, hover over the different lines/boxes to gain information and download the figures as a *.png. The most common way to proceed downstream analysis for single PTM quantification is to calculate relative abundance (RA) as follows^9^:

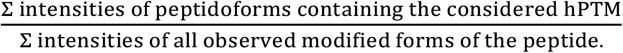

## Code availability

The heatmap shown in **Figure 5** was obtained using an in-house written code (R), available at github (https://github.com/Juliette92/histone_atlas). The PeptidoformViz tool (Shiny App) is available at github (https://github.com/ndmeulem/PeptidoformViz).

## Acknowledgements

This research was funded by mandates from the Research Foundation Flanders (FWO) awarded to BVP (11B4518N), LCo (1SF2622N), ND (1S77220N), SV (3S031319) and MD(12E9716N), the Ghent University Research Fund (BOF), and LM acknowledges funding from the European Union’s Horizon 2020 Programme (H2020-INFRAIA-2018-1) (823839), from the Research Foundation Flanders (FWO) (G028821N), and from Ghent University Concerted Research Action (BOF21/GOA/033). We thank Ian Morns for enabling project licensing in Progenesis and Kelly Giles and Gabrielle Lever from Technology Networks for providing us with the recording of the Progenesis QIP seminar. We also thank Panagiotis Ntziachristos for proofreading.

## Author contributions

L.P.: histone extraction, gel electrophoresis, propionylation, data analysis and validation, writing

B.V.P.: histone extraction, propionylation, data acquisition, data analysis and validation, writing

L.Co.: data analysis and validation, writing

N.D.: heatmap code, PeptidoformViz tool, writing

S.V.: propionylation, feature editing

B.L.: cell culture

S.D.: gel electrophoresis

J.R.: heatmap code

L.Cl.: supervision PeptidoformViz tool

L.M.: supervision PeptidoformViz tool

D.D.: co-supervision

P.V.V.: supervision

M.D.: writing, supervision

## References

1. Grishina, O. et al. DECIDER: prospective randomized multicenter phase II trial of low-dose decitabine (DAC) administered alone or in combination with the histone deacetylase inhibitor valproic acid (VPA) and all-trans retinoic acid (ATRA) in patients >60 years with acute my. BMC Cancer 15, 430 (2015).

2. Dunn, J. & Rao, S. Epigenetics and immunotherapy: The current state of play. Mol. Immunol. 87, 227–239 (2017).

3. Dell’Aversana, C., Lepore, I. & Altucci, L. HDAC modulation and cell death in the clinic. Exp. Cell Res. 318, 1229–1244 (2012).

4. Gonzalez-Lugo, J. D., Chakraborty, S., Verma, A. & Shastri, A. The evolution of epigenetic therapy in myelodysplastic syndromes and acute myeloid leukemia. Semin. Hematol. 58, 56–65 (2021).

5. Huang, H., Sabari, B. R., Garcia, B. A., Allis, C. D. & Zhao, Y. SnapShot: Histone Modifications. Cell 159, 458–458.e1 (2014).

6. Willems, S. et al. Flagging False Positives Following Untargeted LC-MS Characterization of Histone Post-Translational Modification Combinations. J. Proteome Res. 16, 655–664 (2017).

7. Janssen, K. A., Sidoli, S. & Garcia, B. A. Recent Achievements in Characterizing the Histone Code and Approaches to Integrating Epigenomics and Systems Biology. Methods Enzymol. 586, 359–378 (2017).

8. Bian, Y. et al. Robust, reproducible and quantitative analysis of thousands of proteomes by micro-flow LC-MS/MS. Nat. Commun. 11, 157 (2020).

9. De Clerck, L. et al. hSWATH: Unlocking SWATH’s Full Potential for an Untargeted Histone Perspective. J. Proteome Res. 18, 3840–3849 (2019).

10. Errington, T. M., Denis, A., Perfito, N., Iorns, E. & Nosek, B. A. Challenges for assessing replicability in preclinical cancer biology. Elife 10, (2021).

11. Bereman, M. S. et al. An Automated Pipeline to Monitor System Performance in Liquid Chromatography-Tandem Mass Spectrometry Proteomic Experiments. J. Proteome Res. 15, 4763–4769 (2016).

12. De Clerck, L. et al. Untargeted histone profiling during naive conversion uncovers conserved modification markers between mouse and human. Sci. Rep. 9, 17240 (2019).

13. Govaert, E. et al. Extracting histones for the specific purpose of label-free MS. Proteomics 16, 2937–2944 (2016).

14. Meert, P., Govaert, E., Scheerlinck, E., Dhaenens, M. & Deforce, D. Pitfalls in histone propionylation during bottom-up mass spectrometry analysis. Proteomics 15, 2966–2971 (2015).

15. Verhelst, S. et al. Comprehensive histone epigenetics: A mass spectrometry based screening assay to measure epigenetic toxicity. MethodsX 7, 101055 (2020).

16. van Mierlo, G. et al. Integrative Proteomic Profiling Reveals PRC2-Dependent Epigenetic Crosstalk Maintains Ground-State Pluripotency. Cell Stem Cell (2019) doi:10.1016/j.stem.2018.10.017.

17. Deutsch, E. W. et al. The ProteomeXchange consortium in 2020: enabling ‘big data’ approaches in proteomics. Nucleic Acids Res. 48, D1145–D1152 (2020).

18. Van Puyvelde, B. An interactive mass spectrometry atlas of histone posttranslational modifications in T-cell acute leukemia. Panorama Public https://doi.org/10.6069/j26v-sp56 (2022).

19. Dai, C. et al. A proteomics sample metadata representation for multiomics integration and big data analysis. Nat. Commun. 12, 5854 (2021).

20. Roy, U. & Raghavan, S. C. Deleterious point mutations in T-cell acute lymphoblastic leukemia: Mechanistic insights into leukemogenesis. Int. J. cancer 149, 1210–1220 (2021).

21. Schäfer, V. et al. EZH2 mutations and promoter hypermethylation in childhood acute lymphoblastic leukemia. J. Cancer Res. Clin. Oncol. 142, 1641–1650 (2016).

22. Simon, C. et al. A key role for EZH2 and associated genes in mouse and human adult T-cell acute leukemia. Genes Dev. 26, 651–656 (2012).

23. Verhelst, S. et al. A large scale mass spectrometry-based histone screening for assessing epigenetic developmental toxicity. Sci. Rep. 12, 1256 (2022).

24. Goeminne, L. J. E., Gevaert, K. & Clement, L. Peptide-level Robust Ridge Regression Improves Estimation, Sensitivity, and Specificity in Data-dependent Quantitative Label-free Shotgun Proteomics. Mol. Cell. Proteomics 15, 657–668 (2016).

25. R Core Team. R: A language and environment for statistical computing. https://www.r-project.org/ (2017).

26. RStudio Team. RStudio: Integrated Development Environment for R. http://www.rstudio.com/ (2021).

27. Callister, S. J. et al. Normalization approaches for removing systematic biases associated with mass spectrometry and label-free proteomics. J. Proteome Res. 5, 277–286 (2006).

