## Supplementary File 1 for "An interactive mass spectrometry atlas of histone posttranslational modifications in T-cell acute leukemia"

**Randomization scheme.**
Due to the high number of samples, samples were randomized into multiple batches for histone extraction (letter A to C) and propionylation (pink or purple).
When the number of histones extracted from a sample was too low,
an extra histone extraction was performed in either batch D or E.

Histone extraction: batch A → E
Propionylation steps: batch 1 or batch 2

| | REPLICATE | | | | | |
|---|---|---|---|---|---|---|
| **CELL LINES** | 1 | 2 | 3 | 4 | 5 | 6 |
| ALL-SIL | A > E | B > E | C | A | B | C |
| CCRF-CEM | C | A > E | B | C | A > D | B |
| CUTTL-1 | A | B > D | C | A | B | C |
| DND-41 | B > D | C > E | A > D | B | C | A > D |
| HPB-ALL | C | A | B | C | A | B |
| HSB-2 | A | B | C | A | B | C |
| JURKAT | A | B | C | A | B | C |
| KARPAS-45 | C | A | B | C | A | B |
| KARPAS-45 JC | A | B > E | C | A | B | C |
| KE-37 | A | B | C | A | B > D | C |
| KOPTK-1 | B > E | C | A > E | B | C | A > D |
| LOUCY | C | A | B > E | C | A | B |
| MOLT-4 | A | B | C | A | B | C |
| MOLT-16 | B | C | A | B | C | A |
| P12-ICHIKAWA | C | A > E | B | D | A | B |
| PEER | C | A | B | C | A | B |
| PER-117 | C | A > D | B > D | C | A | B |
| PF-382 | B > E | C > D | A > D | B | D | A > D |
| RPMI-8402 | B | C | A | B | C | A |
| SUPT-11 | B | C | A | B | C | A |
| TALL-1 | B | C | A | B | C | A |
