## Supplementary File 2 for "An interactive mass spectrometry atlas of histone posttranslational modifications in T-cell acute leukemia"

### Materials and methods

**Western blot.** Histones from 6 x $10^6$ cells were extracted via the DA extraction method. After extraction, the histone concentration was measured via the BSA protein assay kit. An equal number of histones for each cell line was further prepped for western blot. 5x Laemmli buffer with β-mercaptoethanol was added to the samples in a 1:4 ratio and incubated for 10 minutes at 95°C for denaturation. To perform gel electrophoresis, 20 µl of the samples was loaded on two 9-16% Mini-PROTEAN TGX Precast Protein Gels (Bio-Rad: 456-1103) and 4 µl of the Page Ruler Plus Prestained protein ladder (Perbio: 26620). To identify H3K27me3, the anti-H3K27me3 antibody (07-449; Millipore) was added to one blot (1/1000 in 5% milk/TBST) and incubated overnight. To normalize against the H3 level, the H3 antibody (ab1791; Abcam) was added to the other blot (1/1000 in 5% milk/TBST). After addition of anti-rabbit antibody, both blots were visualized on the Amersham Imager 680 (GE healthcare) via SuperSignal West Dura Extended Duration Substrate.

**Flow cytometry.** A 0,3 million cell pellet of each cell line was resuspended in 100 µl wash buffer (PBS with 2% BSA) before the cells were stained using the Live/Dead Dye efluor 506 (Invitrogen: 65-0866-14). After 30 minutes incubation in the dark at 4°C, the cells were washed with PBS and centrifuged for 5 minutes at 1500rpm at 4°C and the supernatant was drained. Next, the cells were fixated and permeabilized (Invitrogen eBioscience FOXP3/Transcription) to open the membranes for intracellular staining by incubating the cells for 30 minutes at room temperature with 0.5 ml Fix/Perm. After washing the pellets twice with Perm/milliQ water (1/10), the cells were resuspended in 300µl wash buffer and split into three tubes. For each cell line, one tube was stained with H3K27me3 Ab (Millipore; 07-449), another with the H3 Ab (Abcam; ab1791), and a third with only the secondary Ab as negative control to detect background fluorescence and to know the threshold. After 30 minutes incubation in the dark at 4°C, pellets were washed twice before the cells were resuspended in 100µl PBS. Next, the secondary antibody (Thermofisher; AF488) was added and the cells were incubated in the dark at 4°C for 30 minutes. Pellets were washed twice before the cells were resuspended in 150µl PBS and the signal was measured on the BD LSR II (Biosciences). The data was analyzed using FlowJo. Four gating selections were made: SSC vs FSC to deselect debris, SSC-H vs SSC-A to select singlets, AmCyan vs FSC to select living cells, histogram of FITC to visualize the H3 and H3K27me3 level. The measured level of H3K27me3 was normalized against the measured level of H3.

**Mass spectrometry on TripleTOF 5600.** A TripleTOF 5600 mass spectrometer (Sciex, Concord, Ontario, Canada) fitted with a Duospray ion source operating in positive ion mode, was coupled to an Eksigent NanoLC425 HPLC system (Eksigent, Dublin, CA). A volume of each sample, corresponding to 2µg histones, was loaded at 5µL/min with 0.1% Trifluoroacetic acid (TFA) in water and trapped on a YMC TriArt C18 guard column (id 500µm, length 5mm, particle size 3µm) for 5 minutes. Peptides were separated on a microLC YMC TriArt C18 column (id 300 µm, length 15 cm, particle size 3 µm) maintained at 55°C at a flow rate of 5µL/min by means of trap-elute injection. Mobile phase A consisted of UPLC-grade water with 0.1% (v/v) FA and 3% (v/v) DMSO, and mobile phase B consisted of UPLC-grade ACN with 0.1% (v/v) FA. Peptide elution was performed at 5µL/min using the following gradient: i) 3% to 45% mobile phase B in 60 min, ii) ramp to 80% mobile phase B in 5 min. The washing step at 80% mobile phase B lasted 5 min and was followed by an equilibration step at 3% mobile phase B (starting conditions) for 5 min. Ion source parameters were set to 5.5 kV for the ion spray voltage, 30 psi for the curtain gas, 13 psi for the nebulizer gas and 80°C as source temperature.

**DDA**

For DDA (a cycle time of 2.3 s), MS1 spectra were collected between 400-1250 m/z for 250 ms. The 10 most intense precursors ions with charge states 2-5 that exceeded 300 counts per second were selected for fragmentation, and the corresponding fragmentation MS2 spectra were collected between 65-2000 m/z for 200 ms. After the fragmentation event, the precursor ions were dynamically excluded from reselection for 10 s.

**SWATH**

For SWATH (a cycle time of 3 s), a 60 variable window acquisition scheme was used for all samples as described by De Clerck et al. Briefly, SWATH MS2 spectra were collected in high-sensitivity mode from 350-1250 m/z, for 48 ms. Before each SWATH MS cycle an additional MS1 survey scan in high sensitivity mode from 350-1250 m/z was recorded for 96 ms.

### <u>Results</u>

Flow cytometry normalization calculations.

| | **H3K27me3** | **H3** | **Normalized H3K27me3 = H3K27me3/H3** |
|---|---|---|---|
| HSB-2 | 10429 | 7595 | 1,37314 |
| PEER | 14288 | 8882 | 1,608647 |
| JURKAT | 15356 | 10791 | 1,423038 |
| MOLT-16 | 9119 | 5124 | 1,779664 |
| TALL-1 | 15414 | 15372 | 1,002732 |
| RPMI-8402 | 17741 | 16516 | 1,074171 |
| LOUCY | 6621 | 9333 | 0,709418 |
| HPB-ALL | 9276 | 11920 | 0,778188 |
