## Supplementary File 3 for "An interactive mass spectrometry atlas of histone posttranslational modifications in T-cell acute leukemia"

**Supplementary File 6**

| CELL LINE | PLOIDY STATE | |
|---|---|---|
| ALL-SIL | hypertetraploid | 90-95<4n>XX/XXYY, +6, +8, +8, t(1;13)(p32;q32)x2, del(6)(q25)x2, del(9)(?p23p24)x2, t(10;14)(q24;q11.2)x2, add(17)(p11)x2 - sideline with idem, -20, -20 - carries t(10;14) effecting TLX1-TRD (HOX11-TCRD) juxtaposition |
| CCRF-CEM | near-tetraploid | extensive subclonal variation - 90(88-101)<4n>XX, -X, -X, +20, +20, t(8;9)(p11;p24)x2, der(9)del(9)(p21-22)del(9)(q11q13-21)x2 - sideline with +5, +21, add(13)(q3?3), del(16)(q12) |
| CUTTL-1 | hyperdiploid | [50, XY, t(7;9),+16,+18,+20,+21] with a characteristic t(7;9)(q34;q34) translocation |
| DND-41 | near-tetraploid | 12% polyploidy - 92(86-98)<4n>XXYY, -9, -15, +20, +mar |
| HPB-ALL | pseudodiploid | 8% polyploidy - 46<2n>XY, der(1)t(1;16)(q22;p11-12)add(16)(p13), del(2)(p24), del(3)(p11), der(5)t(1;5)(q22-24;q32-33), r(16)(?p12?q12) - sideline with -3 instead of del(3) |
| HSB-2 | pseudodiploid | 4% polyploidy - 46(42-46)<2n>XY, t(1;7)(p34;q35), carries t(1;7) effecting LCK-TRB (LCK-TCRB) fusion |
| JURKAT | hypotetraploid | 7.8% polyploidy - 87(78-91)<4n>XX, -Y, -Y, -5, -16, -17, -22, add(2)(p21)/del(2)(p23)x2 - sideline with additional der(5)t(5;10)(q11;p15), del(9)(p11) |
| KARPAS-45 | hypotetraploid | 84, -Y, -Y, -2, -3, -4, +6, -9, -13, -14, +19, -20, -21, t(X;11)(q13;q23.3), der(X)t(X;11)(q13;q23.3), t(1;5)(q25;q13.1)x2, del(4)(q21.1q31.1),der(11)t(14;11)(11;X)(q11;p13)(q23.3;q13) |
| KARPAS-45 JC (KARPAS-45 cell line deliverd by the lab of Jan Cools) | hypotetraploid | 84, -Y, -Y, -2, -3, -4, +6, -9, -13, -14, +19, -20, -21, t(X;11)(q13;q23.3), der(X)t(X;11)(q13;q23.3), t(1;5)(q25;q13.1)x2, del(4)(q21.1q31.1),der(11)t(14;11)(11;X)(q11;p13)(q23.3;q13) |
| KE-37 | hypotetraploid | 91(86-92)<4n>XXYY, +8, -14, -14, t(7;12)(q32-33;p12-13)x2, der(8)t(8;14)(q24;q11)x4, der(14)t(8;14)(q24;q11)x2 - resembles published karyotype - carries 4 copies of der(8)t(8;14) associated with T-ALL - normal ch 14 present |
| KOPTK-1 | / | 95, XXY, -Y, +8, -11, -11, +12, +13, -14, -14, -15, -20, +21, +mar, del(21)(q22), +der(11)t(11;14)(p13;q11.2)x2, +der(17)t(17;?)(q21;?)x2 |
| LOUCY | hypodiploid | 6% polyploidy - 45<2n>X, -X, del(1)(p3?2p3?4), del(5)(q14-15q34-35), t(16;20)(p1?1;q1?3) - sideline with del(6)(q23) - 1p32, 1p34 and 6q- recurrent in T-ALL |
| MOLT-4 | hypertetraploid | 89-99<4n>XXYY, +4, +7, +8, +20, +20, del(6)(q16)x2, der(7)t(7;7)(p15;q11)x2 |

| | | |
|---|---|---|
| MOLT-16 | near-diploid | 46(43-47)<2n>XX, t(3;11)(p21;p13), t(8;14)(q24;q11) - t(8;14)(q24;q11) and 11p13 breakpoint are associated with T-ALL |
| P12-ICHIKAWA | hypotetraploid | 1.6% polyploidy - 82-84<4n>XX, -Y, -Y, -2, +6, -9, -9, -10, -14, +20, -21, -21, -22, -22, +mar, der(2;9)(p10;q10), del(4)(q25), der(10;22)(q10;q10), add(19)(q13) - sideline with N4, add(8)(p1?) |
| PEER | pseudodiploid | 7% polyploidy - 46(42-47)<2n>XX, der(4)?dup ins(4;4)(?p11;?q21q25), del(5)(q22q31), del(6)(q13q22), del(9)(p11p22), del(9)(q22) - resembles published karyotype - 5q-, 6q-, 9p- and 9q- associated with hematological malignancy |
| PER-117 | near-diploid | 46, XY, -16, +18, t(1;9;11)(p13;p22;p11) |
| PF-382 | near-diploid | 9% near-tetraploid and 15% polyploid sidelines - 45/46(43-47)<2n>X/XX, -10, +14 - sideline with +add(?15)(p11), add(1)(p32) |
| RPMI-8402 | hypotetraploid | 2% polyploidy - 90(79-91)<4n>XXX, -X, +3, +3, -10, -13, -14, +15, -18, -20, +2mar, dup(4)(q13q23)x2, del(6)(q14q22)x2, t(11;14)(p15;q11)x2, add(15)(p13) - sideline with der(1)t(1;9)(p35/36;q11), add(13)(q34) - carries t(11;14) with LMO1-TRD (LMO1-TCRD) rearrangement and cryptic del(1)(p32) effecting STIL-TAL1 (SIL-TAL1) fusion |
| SUPT-11 | hypodiploid | 5% polyploidy; 38(36-39)<2n>XX, -Y, -10, -13, -15, -16, -17, -21, -22, del(X)(q11?q23), add(X)(q23), der(1)del(1)(p32)t(1;4)(q32;q32.2), der(1)t(1;7)(q32;q32), der(2)t(2;4)(p23;p15 or q35), der(3)t(1;3)(p33;q28), der(4)t(1;4)(p32;q32.2), dup(4)(p11p14), der(5)t(5;9)(p15;q11.2), der(6)t(1;6)(p3?2;p2?4), add(6)(q27), der(7)t(7;15)(q31.2;q21.2), add(8)(q24.33), der(8)t(8;16)(p12;p13), del(9)(q21q23), der(12)t(8;12)(q24;p13)t(12;16)(q24;23), t(14;14)(q11.2;q32.1), der(15)t(1;15)(q31;q26), der(19)t(19;21)(p13;q21), der(20)t(17;20)(p11;q13), der(21)t(12;21)(?q24;q12), add(22)(q13) |
| TALL-1 | hypertetraploid | - 90-102<4n>XXY/XXYY, -3, -9, -12, +13, +14, +14, -15, +20, +20, +21, +21, +2-4mar, der(X;1)(p10;p10)/add(1)(q11), add(1)(p11), add(5)(p1?5), del(2)(p22p24), del(12)(q24.1), dup(21)(q11qter)x3-4 |

<u>Used sources</u>

DSMZ - German Collection of Microorganisms and Cell Cultures GmbH

The leukemia-lymphoma cell line facts book by Hans G. Drexler, ISBN 0-12-221970-8, 2001 by ACADEMIC PRESS
